# Impaired Suppression of Plasma Lipid Extraction and its Partitioning Away from Muscle by Insulin in Humans with Obesity

**DOI:** 10.1101/2024.06.11.598550

**Authors:** Christos S. Katsanos, Lee Tran, Nyssa Hoffman, Lori R. Roust, Eleanna De Filippis, Lawrence J. Mandarino, Kailin Johnsson, Marek Belohlavek, Matthew R. Buras

## Abstract

**Context:** Humans with obesity and insulin resistance exhibit lipid accumulation in skeletal muscle, but the underlying biological mechanisms responsible for the accumulation of lipid in the muscle of these individuals remain unknown.

**Objective:** We investigated how plasma insulin modulates the extraction of circulating triglycerides (TGs) and non-esterified fatty acids (NEFAs) from ingested and endogenous origin in the muscle of lean, insulin-sensitive humans (Lean-IS) and contrasted these responses to those in humans with obesity and insulin resistance (Obese-IR).

**Methods:** The studies were performed in a postprandial state associated with steady-state plasma TG concentrations. The arterio-venous blood sampling technique was employed to determine the extraction of circulating lipids across the forearm muscle before and after insulin infusion. We distinguished kinetics of TGs and NEFAs from ingested origin from those from endogenous origin across muscle by incorporating stable isotope-labeled triolein in the ingested fat.

**Results:** Insulin infusion rapidly suppressed the extraction of plasma TGs from endogenous, but not ingested, origin in the muscle of the Lean-IS, but this response was absent in the muscle of the Obese-IR. Furthermore, in the muscle of the Lean-IS, insulin infusion decreased the extraction of circulating NEFAs from both ingested and endogenous origin; however, this response was absent for NEFAs from ingested origin in the muscle of the Obese-IR subjects.

**Conclusions:** Partitioning of circulating lipids away from the skeletal muscle when plasma insulin increases during the postprandial period is impaired in humans with obesity and insulin resistance.

---

Humans with obesity and insulin resistance display accumulation of lipids in skeletal muscle (1,2,3). An important factor in the equation determining lipid accumulation in muscle is the disposal of circulating lipids into the muscle, such as non-esterified fatty acids (NEFAs) and fatty acids released from circulating triglycerides (TGs). Although elevated circulating NEFAs in the fasting-state have been proposed to drive lipid accumulation into the muscle of humans with obesity and insulin resistance (4,5), several lines of evidence argue against this notion (6,7,8). Circulating TGs constitute a source of fatty acids taken by the muscle, and current evidence shows that muscle takes up a substantial portion, if not all, of the fatty acids liberated following extraction of plasma TGs (9,10). This response is of particular importance in the postprandial period (i.e., fed state) because not only TGs from endogenous origin, but also TGs from ingested origin are present in the circulation. Postprandial lipid metabolism is complex, and it has been a challenge to delineate between favorable and unfavorable responses, particularly in regards to how ingested fat is handled in tissues during the postprandial period (11).

Humans with insulin resistance have increased uptake of plasma TGs by the muscle during the postprandial period (12,13). A key step for to the uptake of fatty acids liberated from circulating TGs is the extraction of the circulating TGs by the action of endothelial lipoprotein lipase (LPL) in muscle capillaries. The increase in plasma insulin is a key hormonal response during the postprandial period. In healthy individuals, insulin infusion decreases the activity of LPL in muscle (14). Moreover, insulin sensitivity correlates directly with the insulin-mediated decrease in muscle LPL activity (14), suggesting that the physiological effect of plasma insulin on suppressing the extraction of plasma TGs in muscle would be less evident in the presence of insulin resistance. Further, it is likely that plasma insulin suppresses the extraction of TGs from endogenous origin, as there is a preferential extraction of circulating TGs from ingested origin during the postprandial period (10).

We employed a carefully-designed experimental protocol to determine how acute increase in plasma insulin affects the fractional extraction of plasma TGs in muscle of healthy humans and humans with obesity and insulin resistance. Study participants ingested fat labeled with a stable isotope tracer to discern the specific effects of plasma insulin on regulating the extraction of lipids from ingested origin. The protocol was chosen to avoid recycling of ingested lipid in the endogenous plasma TG pool, and because it takes at least three hours after a meal for fat from that meal to appear in the endogenous TG pool (10,15). We hypothesized that acute hyperinsulinemia immediately suppresses the fractional extraction of plasma TGs in the muscle of healthy, insulin-sensitive humans, but not in humans with obesity and insulin resistance.

## Materials and Methods

### Study design

Seventeen volunteers, ages 18 to 40 years, took part in a study to quantify the fractional extraction of circulating lipids across the forearm muscle. To quantify the extraction of lipids derived specifically from ingested fat, subjects ingested whipping cream enriched with ^13^C- triolein. We determined muscle extraction of both ingested and endogenous plasma TGs and NEFAs, along with that of plasma glucose, at fasting plasma insulin concentrations (Basal Study Period) and during increased plasma insulin concentrations (Insulin Study Period; Figure 1). The arterio-venous (A-V) sampling technique was employed to determine the extraction of circulating substrates across the forearm muscle. The study was conducted according to Declaration of Helsinki principles and all study procedures were carried out after obtaining approval from the Institutional Review Board at Mayo Clinic (IRB # 12005173). Prior to participation, each study participant gave written, informed consent. The study was registered with ClinicalTrials.gov (NCT01860911).

**Figure 1.**
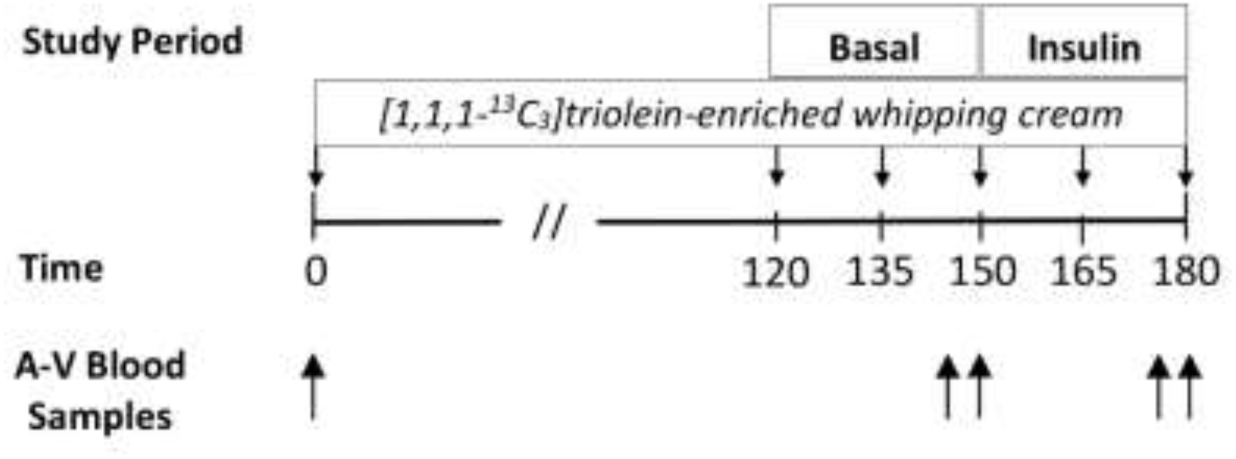
Depiction of the experimental protocol employed to determine the postprandial extraction of plasma triglycerides and non-esterified fatty acids prior to (Basal) and following insulin infusion (Insulin). [1,1,1-^13^C_3_]triolein-enriched whipping cream was ingested at the time points indicated in the form of small boluses to induce steady-state concentrations of ingested and overall plasma triglycerides. Arterio-venous (A-V) blood samples were collected from a radial artery and a vein draining forearm muscle.

### Study participants

Initial screening of interested participants was performed over the phone. Individuals were excluded from the study if they reported diabetes, history of abnormal lipid metabolism, medications, smoking, current participation in a weight-loss regimen, extreme dietary practices, took supplements known to affect the vasculature or lipid metabolism (i.e., cocoa, arginine, protein, fish oil), used anabolic steroids or corticosteroids, or participated in physical activity more than two days per week. Individuals that met the initial criteria were invited for further screening in order to estimate their insulin sensitivity if they had body mass index (BMI) ≤ 25 kg/m^2^ (i.e., lean subjects) or BMI ≥ 32 kg/m^2^ (i.e., subjects with obesity).

### Screening procedures

For the screening visit, participants arrived in the Clinical Studies Infusion Unit (CSIU) at Mayo Clinic in Scottsdale, Arizona, in the morning after an overnight fasting period (i.e., 12 hours). The purpose of the study, the exact experimental procedures, and the associated risks were discussed with each participant prior to obtaining written consent. Screening procedures included medical history, standard physical examination, electrocardiogram (ECG), routine blood tests, and urinalysis. Subjects were excluded from the study if there was evidence of acute illness, metabolic disease such as liver or renal disease, unstable angina, congestive heart failure, plasma TG concentration >2.3 mmol/l, or 2-h plasma glucose > 11.1 mmol/l during an oral glucose tolerance test (OGTT). Insulin sensitivity was estimated from the responses of the plasma glucose and insulin concentrations during the OGTT, and by calculating the Matsuda Insulin Sensitivity Index (ISI) (16). We studied humans with obesity and insulin resistance (Obese-IR) and lean, insulin-sensitive humans (Lean-IS) with Matsuda ISI < 4.5 and > 7.0, respectively. We studied groups of subjects with distinct differences in insulin sensitivity in order to minimize variability in the responses resulting from biological and technical confounding factors. Body composition, including distinct components of body lean and fat tissues (i.e., visceral fat, forearm fat), were evaluated using Dual-energy X-ray absorptiometry (DEXA).

### Infusion study visit

Participants arrived at the CSIU for the infusion study visit at ∼ 7 am after fasting for ∼12 hours. Subjects were asked to maintain their typical diet starting three days prior to the infusion day, and avoid any beverages with alcohol, as well as any form of physical activity during the same period beyond their normal daily physical activities. Compliance with these instructions was verbally verified when participants arrived at the CSIU, and prior to the initiation of the experimental procedures. An intravenous (IV) line was placed in a retrograde fashion deep in an antecubital vein draining the forearm muscle. To ensure that the selected vein drains primarily forearm muscle, oxygen saturation in the blood drawn from this IV line had to be <60% (17); if not, the IV line would be repositioned. Another catheter was placed under local anesthesia in the radial artery of the opposite arm for arterial blood sampling. In addition to the arterial catheter, an additional IV line was inserted in an antecubital vein of the same arm for infusion of insulin. All catheters were kept patent by continuous infusion of saline.

A depiction of the experimental protocol is shown in Figure 1. Background blood samples were obtained at time 0, prior to ingestion of whipping cream containing (per 15 ml) 5 g of fat, 1 g of carbohydrate, and 0 g of protein. Whipping cream was enriched (∼4%) with [1,1,1-^13^C_3_, 99%]triolein (Cambridge Isotope Laboratories, Inc., MA) to differentiate the metabolism of ingested fat in plasma TGs and NEFAs across the forearm muscle. Labeled triolein was incorporated into the whipping cream immediately prior to each experiment by sonication for five mins. After adding a small amount of artificial sweetener, the whipping cream was refrigerated until it was consumed. Subjects ingested a priming bolus of whipping cream [200 mg fat·kg^-1^fat free mass (FFM)], followed by ingestion of smaller boluses starting 2 hours later and every 15 mins thereafter until the end of the infusion study (Figure 1), resulting in an overall fat ingestion of 100 mg fat·kg FFM^-1^·h^-1^ during the course of the experiment. We employed an experimental design that involves ingestion of small boluses of fat in order to avoid increase in the concentration of plasma TGs carrying ingested TG. Lipoprotein particles carrying ingested TG have 50-fold higher affinity for LPL than lipoproteins carrying endogenous TG (18), and in a way that can independently inhibit clearance of lipoproteins carrying endogenous TG by substrate competition for LPL (19). Therefore, by studying the plasma TG metabolism across muscle with no perturbation in the plasma TG concentrations and experimentally manipulating a single variable (i.e., plasma insulin concentration), our experimental approach allowed isolating the specific role of plasma insulin in regulating the extraction of circulating TG across muscle.

To determine the effects of increased plasma insulin concentrations on variables of interest, insulin was infused at a rate of 1.0 mU·kg^-1^·min^-1^ (2.0 mU·kg^-1^·min^-1^ for the first 10 mins) and 0.5 mU·kg^-1^·min^-1^ (1.0 mU·kg^-1^·min^-1^ for the first 10 mins) in the Lean-IS and Obese-IR subjects, respectively. Insulin was infused at a lower rate in the Obese-IR subjects, in order to achieve comparable absolute change in plasma insulin concentrations between the Obese-IR and Lean-IS subjects during the insulin infusion period. To maintain blood glucose concentrations at the basal levels during the insulin infusion, 20% dextrose was infused at a rate that was adjusted based on blood glucose concentration measurements performed every five mins (i.e., hyperinsulinemic-euglycemic clamp procedure). Two blood samples were collected five minutes apart at the end of the Basal and Insulin infusion study periods (Figure 1) to compare the effects of increased plasma insulin on the variables of interest. Arterial and venous blood samples were collected simultaneously from the radial artery line and the line draining the forearm muscle, respectively.

### Blood flow

We measured branchial artery blood flow using Doppler ultrasound in the Basal period and at the end of the 30-min Insulin infusion period in a group of Lean-IS subjects (i.e., first seven Lean-IS subjects), and by following procedures previously described (20). Branchial artery blood flow was measured by evaluating the branchial artery diameter and blood velocity using 2D Doppler ultrasound (CX50, Philips Medical Systems) and a linear array transducer placed over the branchial artery. We recorded two separate 2-minute segments, which were processed at a later time using specialized software. The following equation was used to compute blood flow (expressed as ml·min^-1^): Blood Flow = v·π·(d/2)^2^·60, where v = mean blood velocity (cm·s^-1^), π = 3.14, and d = arterial diameter (cm).

### Analyses of plasma samples

Blood samples for the determination of plasma TGs and NEFAs were collected in tubes containing orlistat, which was added (i.e., 30 ug per ml of blood) to inhibit the in vitro lipolysis of TGs (21). All tubes were kept on ice until they were centrifuged for the collection of plasma samples, which were then stored at -80°C until analyses. An automated blood glucose analyzer (YSI 2300, Yellow Springs, OH) was used to measure the arterial blood glucose concentration during the infusion study in order to obtain feedback on how to adjust the infusion rate of 20% dextrose in response to the insulin infusion. Following the study, concentrations of plasma glucose in arterial and venous samples were determined by averaging measurements obtained in triplicate using the same glucose analyzer. Concentrations of plasma TG in the arterial and venous samples were determined enzymatically using a commercially available assay (Sigma-Aldrich, St. Louis, MO) and by calculating an average concentration based on ten replicates.

TG from ingested origin is carried in plasma chylomicron lipoproteins (TG content: 90%), whereas that from endogenous origin is carried in plasma very-low-density-lipoproteins (VLDL; TG content: 70%) secreted by the liver as well as low-density lipoproteins (LDL; TG content: 10%) resulting from the metabolism of VLDL (22). We distinguished muscle intravascular lipid metabolism originating from ingested fat by analyzing the ^13^C-oleate enrichment in the circulating TGs and NEFAs. Analyses for the determinations of the percentage of oleate within the TG fraction (i.e, oleate-TG), ^13^C enrichment of oleate-TG (i.e., [^13^C]-oleate-TG), concentrations of individual NEFAs (i.e., myristate, palmitate, palmitoleate, palmitelaidate, stearate, oleate, elaidate, linoleate, α-linolenate, arachidonate, eicosapentaenoate, docosahexaenoate), and ^13^C enrichment of oleate-NEFA (i.e., [^13^C]-oleate-NEFA) in the arterial and venous plasma samples were performed at the Mayo Clinic Metabolomics Core. The analyses were performed on a liquid chromatography mass spectrometry (LC-MS) as previously described (23) using an Agilent 6460 triple quadrupole mass spectrometer coupled with a 1290 Infinity liquid chromatography system, allowing for high levels of analytical precision measurements. TGs were isolated by solid phase extraction as described by Bodennec et al. (24). Isolated TGs underwent basic hydrolysis and additional extraction prior to analysis on the LC-MS. Oleate-TG was expressed as a percentage of the sum of 12 individual fatty acids from the same fraction, while [^13^C]-oleate-TG was expressed as tracer-to-tracee ratio (TTR). Concentrations of NEFAs and TTR of [^13^C]-oleate-NEFA in plasma were measured in aliquots that did not go through TG isolation. The concentration of total NEFAs in plasma was calculated as the sum of the measured concentrations of the individual plasma NEFAs indicated above.

Concentrations of amino acids (alanine, arginine, asparagine, glutamine, glycine, isoleucine, leucine, lysine, methionine, phenylalanine, serine, threonine, tryptophan, tyrosine, valine) in the arterial plasma samples were determined using high-performance liquid chromatography as we have described previously (25). The concentrations of plasma insulin (Alpco Diagnostics Cat# 80-INSHU-E01.1, <u>RRID:AB_2801438</u>), apolipoprotein C-II (APOC-II) (Abcam Cat# ab168549, <u>RRID:AB_3107089</u>) and apolipoprotein C-III (APOC-III) (Abcam Cat# ab154131, <u>RRID:AB_3107090</u>) in the arterial plasma samples were measured using commercially available ELISA assays.

### Calculations

The concentration of oleate-TG in arterial and venous plasma samples was calculated by multiplying the measured percentage of oleate in the plasma TGs by the corresponding measured concentration of plasma TGs. The concentration of plasma [^13^C]-oleate-TG was calculated by multiplying the [^13^C]-oleate-TG TTR by the corresponding plasma oleate-TG concentration. The concentration of unlabeled-oleate-TG in plasma was calculated as the difference between the concentration of plasma oleate-TG and that of plasma [^13^C]-oleate-TG. Arterial and venous plasma concentrations of [^13^C]-oleate-NEFA were calculated by multiplying the [^13^C]-oleate-NEFA TTR by the corresponding plasma oleate-NEFA concentration. The concentration of unlabeled-oleate- NEFA was calculated as the difference between the concentration of plasma oleate-NEFA and that of plasma [^13^C]-oleate-NEFA.

We calculated the A-V concentration differences of the plasma metabolites of interest across the muscle to describe metabolite uptake (i.e., positive A-V concentration difference) or release (i.e., negative A-V concentration difference) from the muscle. Fractional extraction of the metabolites of interest across the muscle was calculated as the A-V plasma metabolite concentration difference divided by the arterial plasma metabolite concentration, and expressed as a percentage.

### Statistics

The normality of the data was evaluated using the Shapiro-Wilk test in conjunction with a visual examination of the corresponding Q-Q plots. A logarithmic transformation was applied to datasets that displayed a non-normal distribution prior to performing statistical analysis. Unpaired t-test was used to assess differences in subject characteristics. A two-way (insulin x group) repeated-measures analysis of variance (ANOVA) was employed to examine the main effects of plasma insulin concentrations and obesity/insulin-resistant state on the parameters of interest. Pairwise comparisons were performed using the Bonferroni multiple comparisons test. All tests were two-sided and *P* ≤ 0.05 was considered statistically significant. The APOC-II and APOC-III data were analyzed using a one-sided test, based on the expectation of a specific directional effect (26), as higher concentrations of these apolipoproteins are found alongside insulin resistance and elevated plasma TGs, as observed in our subjects with obesity (27,28). Correlations between variables of interest were evaluated using the Pearson correlation coefficient (*r*). Data are reported as mean ± standard error of the mean (SEM). All statistical analyses were performed using GraphPad Prism Software (version 8.4.3). To enhance the interpretation of our results and to facilitate integration of our data in similar research, we employed a web-based effect size calculator (29) to compute effect size values (i.e., Cohen’s *d*) for the study’s primary end-points.

## Results

Anthropometric and clinical characteristics of the Lean-IS and Obese-IR subjects are presented in Table 1. By design, the two groups differed in BMI and insulin sensitivity.

**Table 1.** Subject characteristics.

|  | Lean - IS | Obese - IR |
| --- | --- | --- |
| n (F/M) | 9 (2/7) | 8 (3/5) |
| Age (years) | 24.6 ± 1.6 | 27.3 ± 3.0 |
| Weight (kg) | 70.6 ± 3.4 | 107.7 ± 7.2*** |
| BMI (kg/m<sup>2</sup>) | 22.4 ± 0.8 | 35.0 ± 2.2*** |
| Lean body mass (kg) | 55.7 ± 2.3 | 65.0 ± 4.7 |
| Forearm lean mass (kg) | 0.99 ± 0.07 | 0.99 ± 0.14 |
| Forearm fat mass (kg) | 0.20 ± 0.03 | 0.56 ± 0.08*** |
| Forearm fat mass (%) | 15.7 ± 2.5 | 35.3 ± 5.5** |
| Body fat mass (%) | 17.5 ± 1.8 | 36.6 ± 3.6*** |
| Android/Gynoid Fat | 0.29 ± 0.03 | 0.62 ± 0.07*** |
| Visceral Fat Mass (g) | 287 ± 68 | 714 ± 135* |
| Fasting plasma glucose (mmol·L<sup>-1</sup>) | 4.6 ± 0.1 | 5.0 ± 0.2* |
| Fasting plasma insulin (μIU·ml<sup>-1</sup>) | 2.9 ± 0.3 | 11.7 ± 1.4*** |
| HOMA-IR | 0.6 ± 0.1 | 2.6 ± 0.3*** |
| Matsuda-ISI | 11.5 ± 1.2 | 3.1 ± 0.3*** |
| HbA1c (mmol·mol<sup>-1</sup>) | 33.5 ± 1.6 | 34.0 ± 1.4 |
| HbA1c (%) | 5.2 ± 0.1 | 5.3 ± 0.1 |
| TSH (mIU·l<sup>-1</sup>) | 1.3 ± 0.3 | 1.5 ± 0.3 |
| Plasma triglycerides (mmol·L<sup>-1</sup>) | 0.8 ± 0.1 | 1.6 ± 0.2** |
| Total plasma cholesterol (mmol·L<sup>-1</sup>) | 3.8 ± 0.2 | 4.1 ± 0.2 |
| Plasma HDL-Cholesterol (mmol·L<sup>-1</sup>) | 1.4 ± 0.1 | 1.0 ± 0.1*** |
| Plasma LDL-Cholesterol (mmol·L<sup>-1</sup>) | 2.0 ± 0.3 | 2.4 ± 0.2 |

Values are mean ± SEM. Lean-IS, lean, insulin sensitive subjects; Obese-IR, subjects with obesity and insulin resistance; BMI, body mass index; Android/Gynoid fat, ratio of android fat mass to gynoid fat mass; HOMA-IR, homeostatic model assessment of insulin-resistance; Matsuda-ISI, Matsuda Insulin Sensitivity Index; HbA1c, glycated hemoglobin; TSH, thyroid-stimulating hormone; HDL, high-density lipoprotein; LDL, low-density lipoprotein.
\* $P \leq 0.05$ , \*\* $P \leq 0.01$ , \*\*\* $P \leq 0.001$ , statistically significantly different from Lean-IS.

### Insulin

Plasma insulin concentrations across the study periods are shown in Table 2. Plasma insulin concentrations were not different between Fasting and Basal study periods in either group, but they were higher (*P* < 0.05) in the Obese-IR than the Lean-IS subjects in both study periods. When compared to the Basal study period, plasma insulin concentrations increased (*P* < 0.05) in both the Lean-IS and the Obese-IR during the insulin infusion. As expected based on our study design to achieve comparable absolute changes between groups in the concentrations of plasma insulin during the Insulin infusion, the delta change from Basal in the plasma insulin concentrations (µIU·ml^-1^) was not statistically significantly different between the Lean-IS and Obese-IR groups (49 ± 3 vs 58 ± 9).

**Table 2.** Concentration of insulin and metabolites in arterial plasma during fasting, the basal period and insulin infusion.

|  | Fasting | Basal | Insulin | ANOVA (*P Value*) |  |  |
| --- | --- | --- | --- | --- | --- | --- |
|  |  |  |  | Interaction | Group | Insulin |
| Insulin ($\mu\text{IU}\cdot\text{ml}^{-1}$) | | | | 0.37 | < 0.001 | < 0.001 |
| Lean-IS | 4.0 ± 0.6 | 4.4 ± 0.7 | 53.3 ± 3.3 | | | |
| Obese-IR | 15.2 ± 1.2 | 16.8 ± 3.1 | 74.9 ± 9.1 | | | |
| Glucose ($\text{mmol}\cdot\text{L}^{-1}$) | | | | 0.42 | 0.45 | 0.82 |
| Lean-IS | 4.4 ± 0.1 | 4.5 ± 0.1 | 4.4 ± 0.1 | | | |
| Obese-IR | 4.6 ± 0.2 | 4.6 ± 0.2 | 4.6 ± 0.1 | | | |
| Total-TG ($\mu\text{mol}\cdot\text{L}^{-1}$) | | | | 0.76 | 0.02 | 0.31 |
| Lean-IS | 585 ± 68 | 642 ± 91 | 589 ± 95 | | | |
| Obese-IR | 1297 ± 280 | 1437 ± 338 | 1384 ± 329 | | | |
| Unlabeled-Oleate-TG ($\mu\text{mol}\cdot\text{L}^{-1}$) | | | | 0.78 | 0.05 | 0.07 |
| Lean-IS | N/A | 218 ± 35 | 198 ± 36 | | | |
| Obese-IR | N/A | 493 ± 135 | 479 ± 132 | | | |
| [$^{13}\text{C}$]-oleate-TG ($\mu\text{mol}\cdot\text{L}^{-1}$) | | | | 0.24 | 0.21 | 0.16 |
| Lean-IS | N/A | 1.5 ± 0.4 | 1.6 ± 0.3 | | | |
| Obese-IR | N/A | 2.8 ± 1.1 | 3.5 ± 1.5 | | | |
| Total-NEFA ($\mu\text{mol}\cdot\text{L}^{-1}$) | | | | 0.17 | 0.07 | <0.001 |
| Lean-IS | 418 ± 72 | 430 ± 36 | 101 ± 12 | | | |
| Obese-IR | 444 ± 53 | 496 ± 48 | 281 ± 38 | | | |
| Unlabeled-oleate-NEFA |  |  |  | 0.10 | 0.02 | <0.001 |
| Lean-IS | N/A | 159 ± 14 | 30 ± 3 | | | |
| Obese-IR | N/A | 174 ± 20 | 89 ± 13 | | | |
| [$^{13}\text{C}$]-oleate-NEFA ($\mu\text{mol}\cdot\text{L}^{-1}$) | | | | 0.01 | 0.55 | 0.42 |
| Lean-IS | N/A | 0.96 ± 0.17 | 0.52 ± 0.07 | | | |
| Obese-IR | N/A | 0.51 ± 0.08 | 0.76 ± 0.22 | | | |

Values are mean ± SEM; Lean-IS, lean, insulin sensitive subjects; Obese-IR, subjects with
obesity and insulin resistance; TG, triglycerides; NEFA, non-esterified fatty acids.

### Glucose

Plasma glucose concentrations are shown in Table 2. There was no statistically significantly difference in plasma glucose concentrations between the Fasting and Basal study periods in either group, nor were there any statistically significantly differences between the groups in any of these study periods. Also, during the insulin infusion, plasma glucose concentrations were not statistically significantly different from Basal in either the Obese-IR or the Lean-IS (Figure 2A).

**Figure 2.**
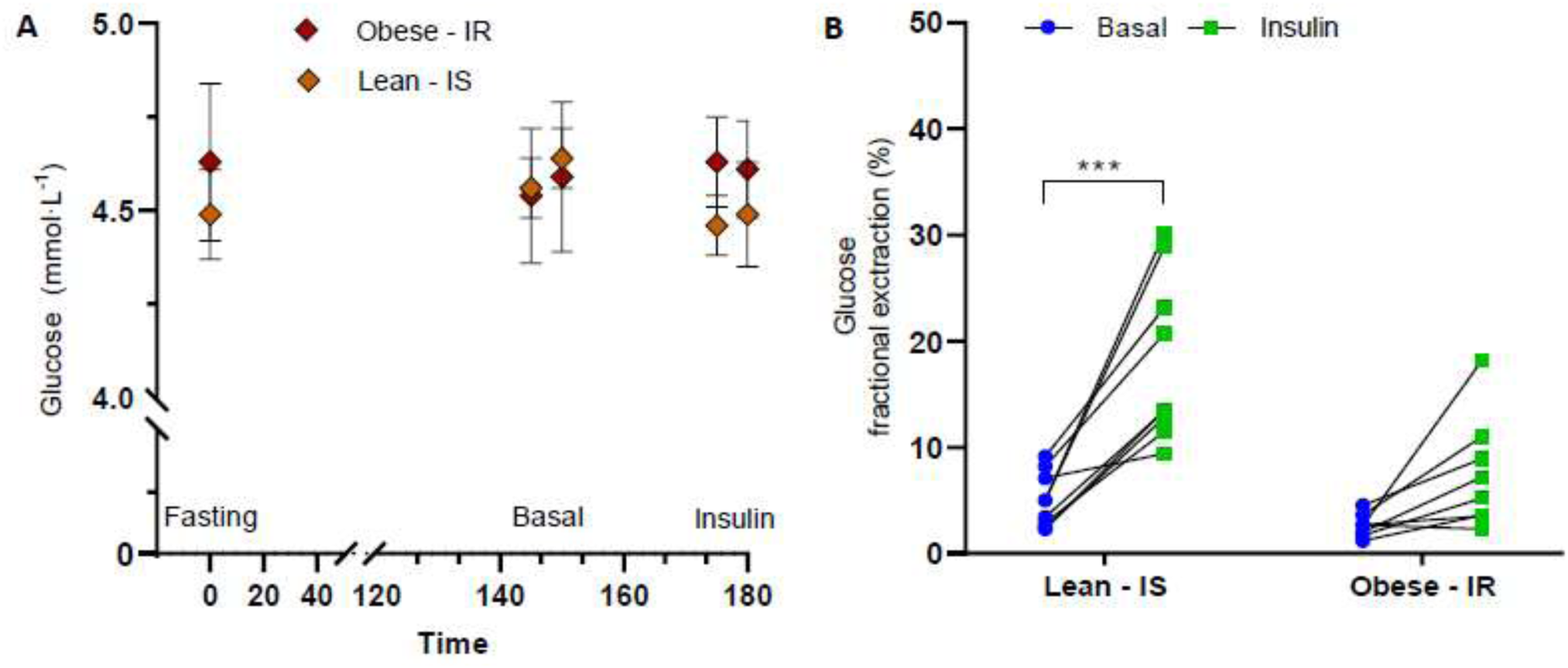
Concentrations of plasma glucose (**A**) and forearm muscle fractional extraction of plasma glucose (**B**) prior to (Basal) and following insulin infusion (Insulin) in lean, insulin sensitive humans (Lean – IS) and humans with obesity and insulin resistance (Obese - IR). Mean ± SEM are shown in (A) and individual data points in (B). Data were analyzed by a two-way ANOVA followed by pairwise comparisons using the Bonferroni multiple comparisons test. \*\*\**P* ≤ 0.001

Plasma glucose fractional extraction (%) across muscle was not statistically significantly different between groups in the Basal period (Lean-IS: 5.0 ± 0.8; Obese-IR: 2.6 ± 0.4), but it increased from Basal in response to the Insulin infusion in the Lean-IS (18.2 ± 2.6; *d* = 2.26), but not in the Obese-IR (7.4 ± 1.9; *d* = 1.28) (Figure 2B). The A-V difference in the concentrations of plasma glucose (Table 3) was not statistically significantly different between groups at Basal, but it increased (*P* < 0.05) in response to the Insulin infusion in the Lean-IS (*d* = 2.09), but not in Obese-IR (*d* = 1.30).

**Table 3.** Arterio-venous differences of metabolites in the basal period and during insulin infusion.

|  | Basal | Insulin | ANOVA (*P value*) |  |  |
| --- | --- | --- | --- | --- | --- |
|  |  |  | Interaction | Group | Insulin |
| Glucose (mmol·L<sup>-1</sup>) |  |  | *0.03* | *0.01* | *<0.001* |
| Lean-IS | 0.23 ± 0.04 | 0.81 ± 0.13 |  |  |  |
| Obese-IR | 0.12 ± 0.02 | 0.35 ± 0.09 |  |  |  |
| Total-TG (μmol·L<sup>-1</sup>) |  |  | *0.59* | *0.46* | *0.03* |
| Lean-IS | 55 ± 15 | 18 ± 6 |  |  |  |
| Obese-IR | 63 ± 27 | 40 ± 14 |  |  |  |
| Unlabeled-oleate-TG (μmol·L<sup>-1</sup>) |  |  | *0.18* | *0.20* | *0.09* |
| Lean-IS | 21 ± 7 | 6 ± 3 |  |  |  |
| Obese-IR | 26 ± 11 | 24 ± 7 |  |  |  |
| <sup>13</sup>C-Oleate-TG (μmol·L<sup>-1</sup>) |  |  | *0.60* | *0.62* | *0.12* |
| Lean-IS | 0.58 ± 0.23 | 0.29 ± 0.16 |  |  |  |
| Obese-IR | 0.96 ± 0.61 | 0.40 ± 0.48 |  |  |  |
| Total-NEFA (μmol·L<sup>-1</sup>) |  |  | *0.04* | *0.50* | *<0.001* |
| Lean-IS | 16.5 ± 17.8 | -45.5 ± 14.6 |  |  |  |
| Obese-IR | 14.8 ± 17.7 | -13.8 ± 14.4 |  |  |  |
| Unlabeled-oleate-NEFA (μmol·L<sup>-1</sup>) |  |  | *0.06* | *0.53* | *<0.001* |
| Lean-IS | 8.7 ± 8.2 | -16.4 ± 6.1 |  |  |  |
| Obese-IR | 8.1 ± 6.4 | -4.3 ± 5.3 |  |  |  |
| <sup>13</sup>C-Oleate-NEFA (μmol·L<sup>-1</sup>) |  |  | *0.04* | *0.53* | *0.05* |
| Lean-IS | 0.18 ± 0.07 | -0.06 ± 0.06 |  |  |  |
| Obese-IR | 0.01 ± 0.06 | 0.01 ± 0.06 |  |  |  |

Values are mean ± SEM; Lean-IS, lean, insulin sensitive subjects; Obese-IR, subjects with obesity and insulin resistance; TG, triglycerides; NEFA, non-esterified fatty acids.

### Amino acids

Plasma total amino acid concentrations (umol·L^-1^) were higher (*P* < 0.05) in the Obese-IR (2652 ± 121) than the Lean-IS (2282 ± 85) at Basal, including the concentrations of the branched chain amino acids (BCAAs; 631 ± 28 vs 487 ± 27). During the insulin infusion, the concentration (umol·L^-1^) of plasma total amino acids was not statistically significantly different from Basal in either the Obese-IR (2560 ± 126) or Lean-IS (2261 ± 68), but that of plasma BCAA was lower (*P* < 0.05) compared to Basal for both the Obese-IR (582 ± 34) and the Lean-IS (463 ± 27).

### Triglycerides

Plasma total TG concentrations were not statistically significantly different between the Fasting, Basal and Insulin infusion periods within either group. Also, the concentrations of neither the unlabeled-oleate-TG or the [^13^C]-oleate-TG were statistically significantly different between the Basal and Insulin infusion periods. In terms of [^13^C]-oleate-TG enrichment, there was a main effect for insulin, but not for group (Table 4); however, there were no statistically significantly differences in either arterial or venous plasma [^13^C]-oleate-TG enrichment in response to the insulin infusion in neither group.

**Table 4.**
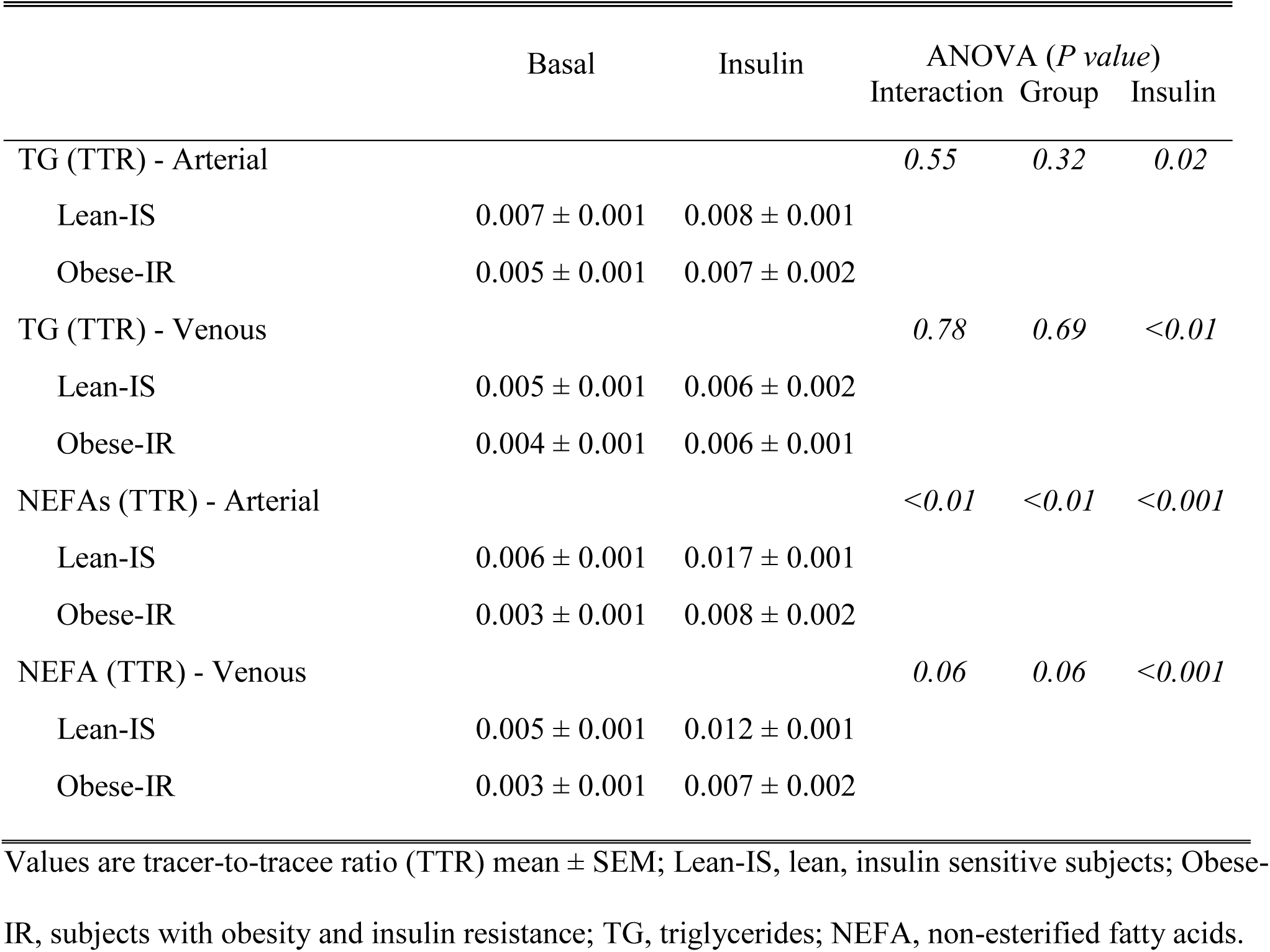
Isotopic enrichment of plasma triglycerides and non-esterified fatty acids with [^13^C]-oleate in arterial and venous plasma in the basal period and during insulin infusion.

Plasma total TG fractional extraction (%) in the Basal period was lower (*P* < 0.05) in the Obese-IR (3.3 ± 0.8) than the Lean-IS (8.8 ± 1.9; *d* = 1.22). In response to the insulin infusion, the fractional extraction (%) of plasma total TG across muscle decreased (*P* < 0.05) when compared to Basal in the Lean-IS (3.4 ± 0.9; *d* = -1.18), but not in the Obese-IR (2.8 ± 0.6; *d* = -0.23). At Basal, the fractional extraction of plasma total TGs in muscle across all subjects correlated statistically significantly and inversely with the concentration of plasma BCAA (*r* = -0.52), but not with that of plasma total amino acids (*r* = -0.42). However, during insulin infusion, there was no statistically significant correlation between the delta changes in the fractional extraction of circulating TGs and the plasma concentrations of BCAA or total amino acids.

The fractional extraction (%) of unlabeled-oleate-TG across the muscle in the Basal Period was lower (*P* < 0.05) in the Obese-IR (4.0 ± 0.9; *d* = 1.15;) than the Lean-IS (9.9 ± 2.3). The fractional extraction of unlabeled-oleate-TG across muscle decreased from Basal in response to the insulin infusion in the Lean-IS (3.3 ± 1.7; *d* = -3.31), but not in the Obese-IR (5.1 ± 1.0; *d* = 1.22) (Figure 3A). In the Basal period, and in contrast to the lower (*P* < 0.05) fractional extraction of unlabeled-oleate-TG in the muscle of the Obese-IR, the fractional extraction (%) of [^13^C]-oleate- TG was not statistically significantly different between the Lean-IS and the Obese-IR (29.5 ± 13.7 vs 22.0 ± 13.6; *d* = 0.18). In response to the insulin infusion, the fractional extraction (%) of plasma [^13^C]-oleate-TG in muscle did not change from Basal in either the Lean-IS (31.2 ± 12.4; *d* = 0.13) or the Obese-IR (-4.7 ± 20.9; *d* = -1.51) (Figure 3B).

**Figure 3.**
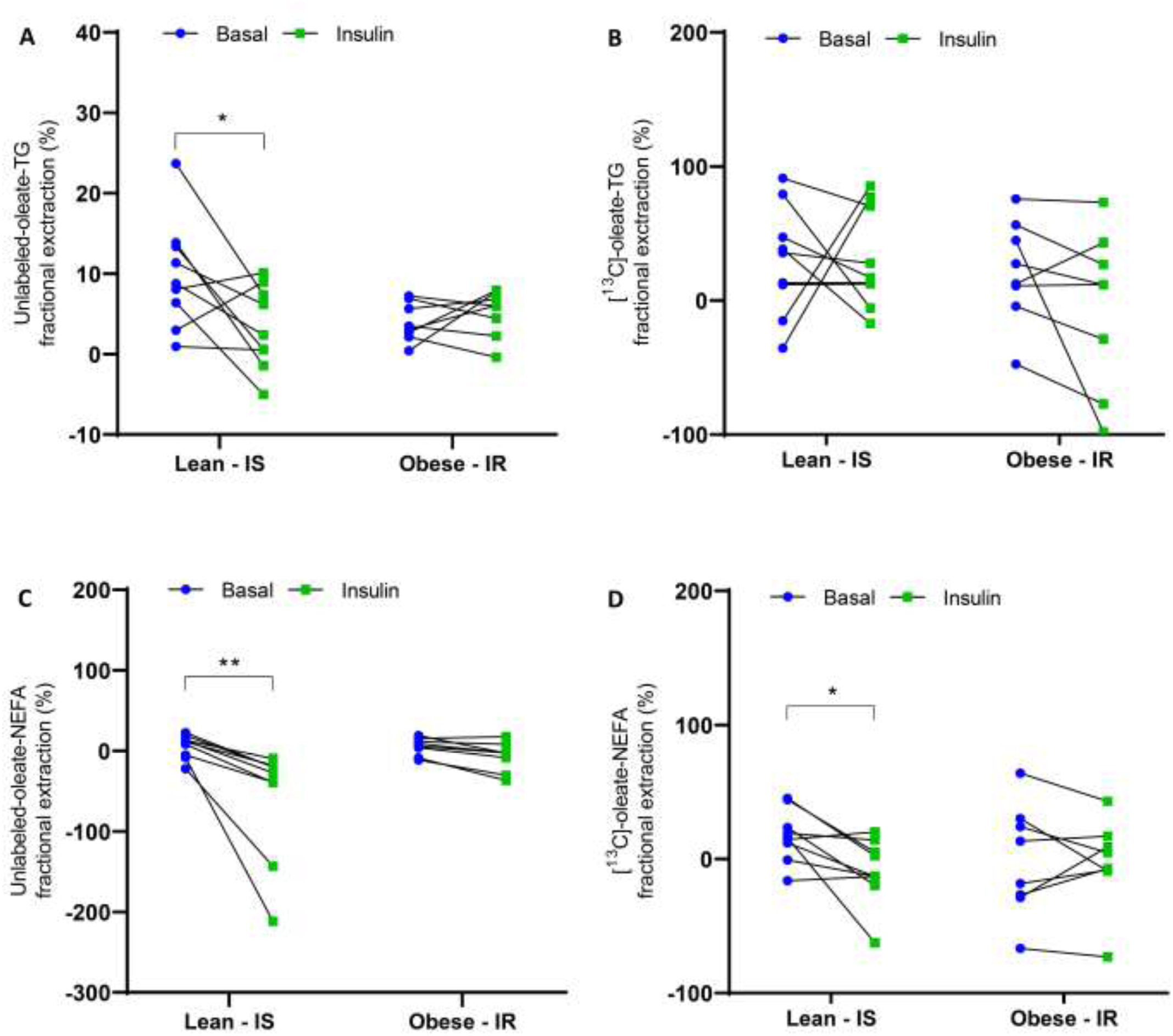
Forearm muscle fractional extraction of plasma endogenously-produced triglycerides (TG) (i.e., Unlabeled-oleate-TG) (**A**) and non-esterified fatty acids (NEFA) (i.e., Unlabeled-oleate-NEFA) (**C**) as well as ingested TG (i.e., [^13^C]-oleate-TG) (**B**) and NEFA (i.e., [^13^C]- oleate-NEFA) (**D**) prior to (Basal) and following insulin infusion (Insulin) in lean, insulin sensitive humans (Lean – IS) and humans with obesity and insulin resistance (Obese - IR). Individual data points are shown. Data were analyzed by a two-way ANOVA followed by pairwise comparisons using the Bonferroni multiple comparisons test. \**P* ≤ 0.05, \*\**P* ≤ 0.01.

At Basal, A-V differences in plasma total TG concentrations were not statistically significantly different between groups (*d* = -0.13). Although not statistically significant, the decrease in A-V differences in plasma total TG concentrations in response to insulin infusion showed a larger effect size in the Lean-IS (*d* = -1.08) compared to the Obese-IR (*d* = -0.39). Similarly, although also not statistically significant, larger effect size was observed for the decrease in A-V differences in plasma concentrations of unlabeled oleate-TG in response to insulin infusion in the Lean-IS (*d* = - 0.93) than the Obese-IR (*d* = -0.07). The decreases in A-V differences in plasma concentrations of [^13^C]-oleate-TG in response to insulin infusion were not statistically significantly different, with effect sizes of *d* = -1.46 and *d* = -1.02 in the Lean-IS and Obese-IR, respectively.

### Non-esterified fatty acids

The concentrations of plasma NEFA in arterial plasma are shown in Table 2. Plasma NEFA concentrations were not statistically significantly different either between the Fasting and Basal study periods within each group, or between the groups during these study periods. Insulin infusion decreased (*P* < 0.05) the concentrations of total plasma NEFA in arterial plasma in both the Lean- IS and the Obese-IR. Moreover, during the insulin infusion, the concentration of total NEFAs in arterial plasma was lower (*P* < 0.05) in the Lean-IS than the Obese-IR. Insulin infusion decreased (*P* < 0.05) the concentration of unlabeled-oleate NEFAs in arterial plasma in both groups, but this decrease was greater (*P* < 0.05) in the Lean-IS. Insulin infusion decreased (*P* <0.05) the concentration of [^13^C]-oleate-NEFAs in the Lean-IS, only. In terms of [^13^C]-oleate-NEFA enrichment, there were main effects for both group and insulin in arterial plasma, but only for insulin in venous plasma (Table 4).

The fractional extraction (%) of total plasma NEFAs in the Basal period was not statistically significantly different between the Lean-IS and Obese-IR (3.6 ± 4.2 vs 3.6 ± 3.8; *d* = 0.01). In response to the insulin infusion, the fractional extraction of total plasma NEFAs became negative (from positive at Basal) in both groups indicating a release of NEFAs from muscle, and this response was statistically significantly different in the Lean-IS (*d* = -4.05), but not in the Obese-IR (*d* = -2.25).

Comparable findings to that of the fractional extraction of plasma total NEFA in the muscle were obtained with respect to the fractional extraction of the unlabeled oleate within the plasma NEFA pool. The fractional extraction (%) of plasma unlabeled-oleate-NEFAs across muscle in the Basal period was nto statistically significantly different between the Obese-IR (5.5 ± 3.8) and Lean-IS (5.6 ± 4.9) subjects (*d* = 0.01). In response to the insulin infusion, the fractional extraction of plasma unlabeled-oleate NEFAs decreased in the Lean-IS (-58.6 ± 23.4; *d* = -3.81) but not in the Obese-IR (-7.2 ± 6.4; *d* = -2.39) (Figure 3C). The fractional extraction (%) of plasma [^13^C]- oleate-NEFA in muscle in the Basal period was not statistically significantly different between the Lean-IS and the Obese-IR (17.4 ± 6.4 vs -1.13 ± 14.7; *d* = 0.58). In response to the insulin infusion, the fractional extraction (%) of plasma [^13^C]-oleate-NEFA in muscle decreased from Basal in the Lean-IS (-8.9 ± 8.1; *d* = -3.59) but not in the Obese-IR (-3.0 ± 11.7; *d* = -0.14) (Figure 3D).

Although the A-V difference in plasma total NEFA concentrations was not statistically significantly different between groups at Basal, the A-V difference decreased (*P* < 0.05) in response to the insulin infusion in both the Lean-IS (*d* = -1.27) and the Obese-IR (*d* = -0.62). Also, the A-V differences in the concentrations of unlabeled-oleate within the plasma NEFA pool decreased (*P* < 0.05) in response to the insulin infusion in both the Lean-IS (*d* = -1.14) and Obese-IR (*d* = -0.73). However, the A-V difference in the concentrations of [^13^C]-oleate within the plasma NEFA pool in response to the insulin infusion decreased (*P* < 0.05) in the Lean-IS (*d* = -1.21) but not in the Obese-IR (*d*= 0.03).

### Blood flow

The branchial artery blood flow (ml·min^-1^) during the insulin infusion was not statistically significantly different from Basal (107 ± 8 vs 102 ± 12; *d* = 0.71).

### Apolipoproteins CII and CIII

The concentrations of plasma APOC-II (mg·L^-1^) in the fasting state were not statistically significantly different between the Lean-IS (31 ± 6) and Obese-IR (23 ± 3) (*d* = 0.52). However, the concentrations of plasma APOC-III (mg·L^-1^) were higher (*P* < 0.05) in the Obese-IR (36 ± 5) than the Lean-IS (26 ± 2.0) (*d* = 0.98).

## Discussion

Our findings show that the postprandial increase in plasma insulin acutely suppresses the fractional extraction of circulating TGs in muscle. However, this response is absent in the muscle of humans with obesity and insulin resistance. As a result, during postprandial hyperinsulinemia, the muscle of individuals with obesity/insulin resistance are exposed to higher amounts of fatty acids released from circulating TGs as they pass through the muscle, a situation compounded by their already elevated plasma TG concentrations of endogenous origin. Additionally, the greater fractional extraction of circulating NEFAs under these conditions in individuals with obesity/insulin resistance suggests increased plasma fatty acid disposal into their muscles. These responses may facilitate lipid accumulation in the muscles of humans with obesity/insulin resistance during the postprandial period. This notion is supported by a recent study conducted under experimental conditions comparable to this study, which found that insulin infusion increased muscle diacylglycerol content in individuals with insulin resistance (30).

The fractional extraction of plasma TGs in muscle is mediated by the activity of intravascular LPL. When assessed *in vitro*, the activity of LPL in muscle is lower in humans with obesity and insulin resistance (31). Our findings confirm this observation *in vivo* by showing lower fractional extraction of circulating TGs in muscle of humans with obesity/insulin resistance. The mechanisms that sustain lower fractional extraction of plasma TGs (i.e., lower LPL activity) in the non-insulin stimulated state in the muscle of humans with obesity/insulin resistance are currently unknown. At Basal, and when evaluated across subjects, we found an inverse correlation between plasma BCAA concentrations and fractional extraction of plasma TGs in muscle. However, a mechanism linking higher plasma BCAA concentrations to lower TG hydrolysis (i.e., lower LPL activity) in muscle remains to be identified. On the other hand, there is evidence for lower LPL gene expression in the muscle of humans with obesity (32). Also, we found that the plasma concentrations of APOC-III were higher in our subjects with obesity/insulin resistance. Because APOC-III directly inhibits LPL from hydrolyzing TGs (33), lower fractional extraction of plasma TGs in the muscle of humans with obesity/insulin resistance at the basal state in our study may be sustained by increased levels of APOC-III in the circulation of these individuals. It is important to emphasize that the physiological significance of any given fractional extraction of plasma TGs in muscle must be evaluated in the context of the prevailing plasma TG concentrations. In our study, the plasma TG concentrations were ∼2.3 times higher in the subjects with obesity/insulin resistance. Because of that, the absolute amount of TGs extracted by the muscle in the basal, non-insulin-stimulated, state would be comparable between subjects with obesity/insulin resistance and lean controls, as demonstrated by comparable A-V differences in the concentrations of plasma TGs.

Our study provides novel evidence that the physiological role of plasma insulin is to acutely (i.e., within minutes) decrease the extraction of plasma TGs in muscle. Because the hyperinsulinemia observed during the postprandial period does not increase the gene expression of LPL in muscle (13), it is expected that the increase in plasma insulin in the present study downregulated the activity, but not the content, of LPL in muscle. In fact, hormones and dietary factors are the primary regulators of the activity of the LPL (34,35). At this time, we can only speculate about the mechanisms by which the activity of LPL in muscle is downregulated by an increase in plasma insulin. One potential mechanism is the stimulation of nitric oxide production in the muscle vasculature by plasma insulin (36). Several lines of evidence show that nitric oxide reduces the activity of LPL (37,38,39), likely through nitration of LPL (40). However, such an LPL-mediated mechanism remains be elucidated *in vivo* in humans, to explain the decrease in the fractional extraction of plasma TGs we observed in muscle in response to acute hyperinsulinemia in our lean/insulin-sensitive subjects.

From a physiological perspective, a decrease in the fractional extraction of plasma TGs in muscle when plasma insulin levels rise - such as during the postprandial period - indicates that TG metabolism is partitioned away from the muscle during the postprandial period in healthy, insulin sensitive humans. However, the absence of a comparable decrease in the fractional extraction of plasma TGs in the muscle of individuals with obesity/insulin resistance by increased plasma insulin, coupled with higher plasma TG concentrations in these individuals, exposes their muscles to greater amounts of fatty acids liberated from circulating TGs as they pass through the muscle during the postprandial period. Given that muscle takes up a large amount, if not all, of the fatty acids liberated following the local extraction of plasma TGs (9,10), the inability to decrease the postprandial fractional extraction of circulating TGs in the muscle of individuals with obesity/insulin resistance can contribute to lipid accumulation in the muscle of these individuals during the postprandial period.

Our findings, which show that the fractional extraction of plasma NEFAs in skeletal muscle is suppressed by the rise in plasma insulin, are consistent with previous research that demonstrated that insulin reduces the fractional extraction of plasma NEFAs in the heart muscle (41). The failure of subjects with obesity/insulin resistance to suppress the fractional extraction of plasma NEFAs from both endogenous and ingested origin during hyperinsulinemia may result from the higher availability of these plasma NEFAs during the insulin infusion period and/or an increased content of CD36 fatty acid transporters, which transport fatty acids into muscle (42), in the sarcolemma of muscle cells of humans with obesity (43). The lack of suppression of fractional extraction of plasma NEFAs during insulin infusion in our subjects with obesity/insulin resistance, combined with their inability to decrease the plasma NEFA concentrations to the same extent as the lean controls, suggests greater NEFA disposal to muscle, likely contributing to lipid accumulation in the muscle of subjects with obesity/insulin resistance during the postprandial period.

## Conclusions

We demonstrate for the first time a direct physiological role of the plasma insulin increase during the postprandial period in partitioning plasma lipids away from skeletal muscle. This is achieved by inhibiting the fractional extraction of plasma TGs from endogenous (but not ingested) origin, and by reducing the fractional extraction of circulating NEFAs from both endogenous and ingested origin. More importantly, we show that these plasma insulin-mediated responses are either absent or reduced in individuals with obesity and insulin resistance. Thus, the capacity of humans with obesity and insulin resistance to redirect circulating lipids away from the skeletal muscle when plasma insulin rises during the postprandial period is impaired.

## Acknowledgments

We gratefully acknowledge the assistance of the subject recruitment team and the nursing staff in the Clinical Studies Infusion Unit at Mayo Clinic in Scottsdale, Arizona for their support in conducting the studies. We also thank the subjects for their participation and commitment to the study procedures.

## Data Availability

Original data generated and analyzed during this study are included in this published article.

